# Predicting Brain Amyloid Positivity from T1 weighted brain MRI and MRI-derived Gray Matter, White Matter and CSF maps using Transfer Learning on 3D CNNs*

**DOI:** 10.1101/2023.02.15.528705

**Authors:** Tamoghna Chattopadhyay, Saket S. Ozarkar, Ketaki Buwa, Sophia I. Thomopoulos, Paul M. Thompson, the Alzheimer’s Disease Neuroimaging Initiative

**Author notes:** All the authors are with the Imaging Genetics Center, Mark and Mary Stevens Neuroimaging and Informatics Institute, Keck School of Medicine, University of Southern California, Marina del Rey, CA, United States.

## Abstract

Abnormal β-amyloid (Aβ) accumulation in the brain is an early indicator of Alzheimer’s disease and practical tests could help identify patients who could respond to treatment, now that promising anti-amyloid drugs are available. Even so, Aβ positivity (Aβ+) is assessed using PET or CSF assays, both highly invasive procedures. Here, we investigate how well Aβ+ can be predicted from T1 weighted brain MRI and gray matter, white matter and cerebrospinal fluid segmentations from T1-weighted brain MRI (T1w), a less invasive alternative. We used 3D convolutional neural networks to predict Aβ+ based on 3D brain MRI data, from 762 elderly subjects (mean age: 75.1 yrs. ±7.6SD; 394F/368M; 459 healthy controls, 67 with MCI and 236 with dementia) scanned as part of the Alzheimer’s Disease Neuroimaging Initiative. We also tested whether the accuracy increases when using transfer learning from the larger UK Biobank dataset. Overall, the 3D CNN predicted Aβ+ with 76% balanced accuracy from T1w scans. The closest performance to this was using white matter maps alone when the model was pre-trained on an age prediction in the UK Biobank. The performance of individual tissue maps was less than the T1w, but transfer learning helped increase the accuracy. Although tests on more diverse data are warranted, deep learned models from standard MRI show initial promise for Aβ+ estimation, before considering more invasive procedures.

**Clinical Relevance:** Early detection of Aβ positivity from less invasive MRI images, could offer a screening test prior to more invasive testing procedures.

## I. Introduction

Alzheimer’s disease (AD) affects over 20 million people worldwide [22]. In the US, one in three elderly people suffer from Alzheimer’s or other forms of dementia. The main cause of the disease is the abnormal accumulation of beta-amyloid protein deposits in the brain, and accumulation of abnormal tau protein in neurons. Positron emission tomography (PET) is generally used with amyloid- and tau-sensitive radioactive tracers to track the pattern of Aβ build-up in the brain.

These PET scans have three major issues - they are expensive, are not widely available and involve the injection of radioactive tracers, which have some risk of adverse effects. The other method to obtain reliable estimates of brain amyloid load is via measurement of amyloid levels in the cerebrospinal fluid (CSF) via spinal tap or lumbar puncture, a highly invasive and painful procedure.

With the approval by the US Food and Drug Administration (FDA) of the drug, lecanemab - an anti-amyloid beta protofibril antibody - for the treatment of mild cognitive impairment or early dementia due to Alzheimer’s disease, there is a growing interest in non invasive testing for Aβ positivity (Aβ+) as a more convenient means to screen patients prior to more invasive testing [26].

Although standard anatomical MRI is not used for detecting amyloid deposition, Aβ build-up leads to brain cell loss which is evident as atrophy on T1-weighted MRI, along with the expansion of the ventricles and widening of the cortical sulci. Islam and Zhang (2020) proposed a Generative Adversarial Network approach to generate synthetic PET images for controls, MCI and AD subjects. We used 3D convolutional neural networks (CNNs) to predict amyloid positivity using 3D CNNs and transfer learning, with 3D T1w brain MRI scans as input. We also tested the model performance by pre-training the model on prior tasks of age and sex prediction from MRI, in 20,000 subjects from the UK Biobank dataset. CNNs are attractive, as they can learn predictive features from raw images without the need for pre-processing. Transfer learning is a type of artificial intelligence/deep learning method that can boost MRI-based AD classification performance. In this technique, some of the network weights are optimized on prior tasks and then ‘frozen’, while others are allowed to be trained on the new task. We also tested how well amyloid positivity (Aβ+) could be predicted from 3D brain MRI with a different approach, still using CNNs, but based on first segmenting the gray matter, white matter and cerebrospinal fluid from T1w brain MRI.. The idea is that amyloid may affect one of the pre-segmented partitions preferentially, making prior tissue classification helpful for the task, and perhaps making training more efficient, as there may be fewer features to learn. We also used similar pre-training techniques from 10,000 subjects from the UK Biobank to evaluate whether it boosted performance on the downstream Aβ+ prediction task. There is some debate about when such pre-training techniques will improve performance on downstream tasks due to differences in the domains of the tasks. Here we tested whether these pre-training techniques improve accuracy. We also examined how strongly the amount of data used in pre-training affects the accuracy in the downstream task.

## II. DATA

The Alzheimer’s Disease Neuroimaging Initiative (ADNI) is a multisite study, launched in 2004 at 58 sites across North America, collecting neuroimaging, clinical and genetic data to better understand biomarkers associated with healthy aging and Alzheimer’s disease. From the ADNI dataset – which is publicly available at adni.loni.usc.edu – we analyzed 3D T1-weighted brain MRI scans from 762 subjects (age: 75.1 years ± 7.6 SD; 394 F/368 M) with a distribution of 459 controls (CN), 67 with mild cognitive impairment (MCI), and 236 with AD. These subjects were selected as they also had available amyloid-sensitive PET scans collected close to the time of the MRI, where the maximum interval between scans was set to 180 days. Subjects who were missing basic clinical information or poor-quality imaging data – such as scans with severe motion, distortion, or ringing – were not included in the final dataset.

The cut-off for amyloid levels, to define Aβ+, was defined by the ADNI Neuroimaging Core based on PET cortical SUVR uptake (denoted as Aβ_1 by ADNI) determined by either mean 18F-florbetapir (Aβ+ defined as >1.11 for cutoff) or florbetaben (Aβ+ defined as >1.20 for cutoff), normalized by using a whole cerebellum region. For the pre-training task, we used 3D T1-weighted (T1w) brain MRI scans from 10,000 subjects (age: 64.59 years ± 7.64 SD; 4,860 F/5,139 M) from the UK Biobank. The T1-weighted brain MRI volumes were pre-processed using a sequence of steps, including nonparametric intensity normalization (N4 bias field correction), ‘skull-stripping’ for brain extraction, registration to a template with 6 degrees of freedom (rigid-body) registration and isometric voxel resampling to 2 mm. Pre-processed images were of size 91×109×91. The T1w images were scaled to take values between 0 and 1 via min-max scaling. The gray matter, white matter and CSF segmentations from the T1w MRI scan were obtained using FreeSurfer [25].

## III. MODEL AND METHODS

After registering the images to a common template, the data was split into independent training, validation and testing sets in the ratio of approximately 70:20:10 (**Table 1**). To augment the training data, we used elastic deformation, a technique often used in medical image processing. We used displacement vectors and a spline interpolation for input image deformation. The 3D CNN architecture (**Figure 1**) consisted of four 3D Convolution layers with a 3×3 filter size, followed by one 3D Convolution layer with a 1×1 filter, and a final Dense layer with a sigmoid activation function. All layers used the ReLu activation function and Instance Normalization. Dropout layers, with a dropout rate of 0.5, and a 3D Average Pooling layer with a 2×2 filter size were added to the 2^nd^, 3^rd^, and 4^th^ layers. Models were trained with a learning rate of 1e-4, and test performance was assessed using balanced accuracy. To deal with overfitting, both L1 and L2 regularizers were used, along with dropouts between layers and early stopping. Hyperparameter tuning was performed by running *k*-fold cross validation.

**Figure 1.**
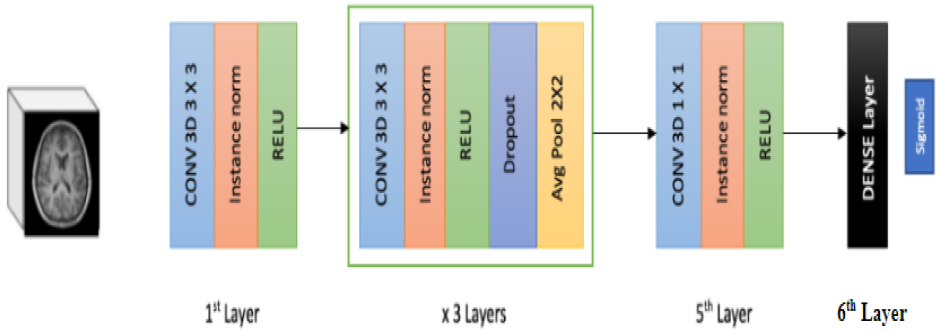
3D CNN Architecture used for training on the ADNI dataset, and for pre-training on the UK Biobank dataset.

**TABLE I.** Distribution of data into independent train, validation and test sets

| Data | N | Age (mean ± SD) | Sex (F/M) | CN/MCI/Dementia |
| --- | --- | --- | --- | --- |
| Training | 530 | 75.3 ± 7.6 | 274/256 | 318/48/164 |
| Validation | 161 | 74.7 ± 7.4 | 83/78 | 97/10/54 |
| Testing | 71 | 74.6 ± 7.5 | 38/33 | 44/9/18 |
| Total | 762 | 75.1 ± 7.6 | 394/368 | 459/67/236 |

For the pre-training task, we used a traditional supervised learning approach based on labeled training data. The initial state of the network was defined by using the below 3D-CNN architecture to predict the sex of the subjects from the T1ws and the Gray Matter, White Matter and CSF maps from the UK Biobank cohort. The 3D-CNN was trained for 40 epochs for each scan with the Adam (with weight decay) optimizer, a learning rate of 1e-4 and a learning rate scheduler. This trained model was fine-tuned to predict Aβ+ using three methods – the model was used with the trained weights; the model’s last two layers were unfrozen, and the model was fine-tuned end to end. In all three methods, the batch size was kept as 6 and the model was trained until the validation loss did not improve for 10 consecutive epochs. We also wanted to understand the effect of the amount of data in the upstream task on the downstream task in pre-training. To this end, the UK Biobank dataset was divided into 8 batches, four of which were of 250 images, and the rest were of 1,000 each while training the model. For T1ws, the UK Biobank dataset was divided into 8 batches of 2000 images each while training the model. Weights for each of these models were stored as the starting weights for downstream tasks. Thus, while fine tuning, the accuracies were calculated for all 8 sets of initial starting weights.

The same method was used for pre-training on an age prediction task (a common benchmark task for CNNs trained on MRIs), i.e., the initial state of the network was defined by using the above 3D CNN architecture to predict the age of the subjects from the T1w and the Gray Matter, White Matter and CSF maps from the UK Biobank cohort. The loss function in this case was the mean square error. Fine-tuning for downstream tasks was the same as in the prior experiments, with the same hyperparameter values. The model was evaluated using metrics that included balanced accuracy and F1 Score using a threshold obtained with the Youden’s Index. Each model was run three times, and the average value was reported.

## IV. RESULTS

The value of Youden’s *J* Index, used to decide the threshold to classify Aβ positivity, was found to be 0.494 when all subjects were considered. A balanced accuracy score of 0.760 with an F1 score of 0.746 was obtained for classification when using the T1w as test data from **Table II.**

**TABLE II.** Comparison of Final Results

| Image Type | Balanced Accuracy |
| --- | --- |
| T1w | 0.760 |
| Gray Matter Map with UK Biobank Age Prediction model used as pre-training | 0.734 |
| White Matter Map with UK Biobank Age Prediction model used as pre-training | 0.739 |
| CSF Map with UK Biobank Sex Classification model used as pre-training | 0.702 |
| Combined SVM model ran on results of CNNs | 0.700 |

Both the pre-training algorithms didn’t have any boosting effect on the classification accuracy. Aβ positivity prediction was performed with around 0.69 balanced accuracy for models pre-trained on sex classification from the UK Biobank dataset. The accuracy was a little less - around 0.66 - when the model was pre-trained on age prediction from the UK Biobank dataset. For the pretraining task of UK Biobank Sex classification, the highest accuracy was 0.531 when all the data was used, and the model had all layers frozen, which was an expected random performance. When the bottom two layers were unfrozen, the highest average balanced accuracy was 0.66 at 12000 images used in pretraining. When the model was trained end-to-end, the average balanced accuracy reached 0.69 at 10000 data points from UK Biobank in the preliminary training task.

When we used the UK Biobank Brain Age Prediction Network without any adaptation (all layers frozen), the performance was at chance level (around 0.457 average balanced accuracy).

When the bottom two layers were unfrozen, the highest accuracy was 0.65 at 12000 images used in pretraining. When the model was trained end-to-end, the accuracy reached 0.678 at 16000 images from UK Biobank in the preliminary training task. Using the values of balanced accuracy and number of data points, we trained a linear regression model to find the slope of the line. Based on our experiment values, we found that in the case of pretraining on age prediction, the slope of the line was 61.843 and in the case of pretraining on sex classification, the slope of the line was 0.821. Positive values on both slopes indicate an increasing curve.

From these pre-training results, four broad conclusions can be made. The first observation was that pre-training on a huge dataset did not boost the accuracy of the downstream task in this scenario. The second observation was that the amount of data used in pre-training did not affect the downstream amyloid positivity prediction task accuracy. The third observation was that models gave almost similar results with minute variations in balance accuracy after 2000 data points being used in the pretraining task. The fourth observation was that pretraining to predict the sex from the T1w MRIs of UK Biobank gave marginally better accuracy on the downstream task of amyloid prediction than pretraining to predict the age from the T1w MRIs.

For gray matter images, the balanced accuracy when the 3D CNN was trained on the ADNI dataset was 0.69. The performance of pretrained models for both age and sex prediction on the UK Biobank data in the upstream task, was similar to performance without any pre-training. The best balanced accuracy was around 0.707 when the sex classification pre-trained model was fine tuned end-to-end. Increasing the amount of data in the upstream task had minimal effect on the balanced accuracy in the downstream task, except for the case of the model which was pre-trained on age prediction on UK Biobank, and then only the bottom two layers were unfrozen for fine tuning. In that case, the balanced accuracy rose numerically to 0.730 when 5,000 subjects were used in the upstream task. To test if there was any statistical gain in accuracy, we fitted a linear regression model to find the slope of the line; the value of the slope was 6.83E-06 with a p-value of 0.263, suggesting no evidence for an increase in accuracy.

For white matter images, the balanced accuracy when the 3D CNN was trained on the ADNI dataset was 0.673. In this case, pretraining the model on upstream tasks such as age prediction and sex classification on UK Biobank dataset actually improved the performance on the downstream task. The best performance was for the model pre-trained on age prediction, with the bottom two layers unfrozen and fine-tuned. The best accuracy was around 0.74 - close to the baseline accuracy for the raw T1ws. We also tested whether increasing data in the upstream task improved performance in the downstream task. The value of the slope in the case of white matter images was 3.84E-02 with a p-value of 0.132. Thus, statistical testing revealed no evidence for performance gain with more pre-training data, at least in the range tested.

For cerebrospinal fluid images, the balanced accuracy when the 3D CNN was trained on the ADNI dataset was 0.671. Perhaps surprisingly, the performance of pretraining the model on the upstream tasks of age prediction and sex classification on the UK Biobank dataset was similar to performance without any pre-training. The best balanced accuracy was around 0.702 when the sex classification pre-trained model was fine-tuned end-to-end. Increase in the amount of data in the upstream task did not have a detectable effect on the balanced accuracy in the downstream task, except for the case of the model which was pre-trained on sex classification on UK Biobank, and then the entire model was fine-tuned.

From these pre-training results, four broad conclusions can be made. First, pre-training on a very large dataset did not boost the accuracy of the downstream task in most scenarios, except in the case of white matter. Second, the amount of data used in pre-training did not affect the downstream amyloid positivity prediction task accuracy, at least in the range tested (very small pre-training datasets would presumably give poor performance). Thirs, models gave comparable results with minute variations in balanced accuracy after 4,000 data points were used in the pretraining task. Finally, pretraining to predict sex from white matter maps in the UK Biobank gave marginally better accuracy on the downstream task of amyloid prediction than pretraining to predict the age from the white matter MRIs. On the contrary, in the case of gray matter and CSF, the performance was better with age prediction as the upstream task. Overall,pre-training on UK Biobank data for age prediction or sex classification did not show substantial improvements in performance, by contrast with training the model directly on the ADNI dataset.

Based on the results from CNNs, we ran a SVM on the outputs, to find the accuracy when we combine the T1w results with various segmented maps. The best accuracy in this case was 0.7 when the SVM had the results from T1w and end to end fine tuned Grey Matter, White Matter and CSF maps as input.

## V. CONCLUSION AND DISCUSSION

Through our experiments, we present results comparing pre-training strategies for amyloid positivity detection from T1ws as well as gray matter, white matter and cerebrospinal fluid segmented images from T1w MRIs. We compared the performance of a 3D Convolutional Neural Network trained from scratch on the data to the performance of models pre-trained on the UK Biobank dataset for two tasks – sex classification and age prediction. We explored three finetuning methodologies – first, where all layers were kept frozen; second, where the bottom two layers were unfrozen; and third, where the entire model is trained end-to-end. We also examined the impact of the amount of data used in pre-training.

Based on our first observation, we can see that the accuracy is greater without the pre-training methodology. This may be because the UK Biobank primarily consists of healthy subjects, whose MRIs may not provide ideal predictive features for amyloid detection in ADNI. Also, though amyloid is not detectable directly with standard anatomical MRI, the 3D CNN architecture could predict amyloid positivity with 76% accuracy. A key question often asked is whether pre-training boosts task performance especially when the amount of training data for the new task is limited. From our experiments, it can be shown that pre-training is not always helpful in improving performance on low data regimes, particularly in tasks where downstream accuracy in predicting is already difficult to achieve and the upstream task has data, which is different, despite being in the same domain.

Increasing the amount of data in upstream task also did not affect the classification performance of the downstream task as seen from Fig 2. There was a plateau effect, where the model did not learn anything new after it reached maximum accuracy from a task. Another observation was the increase in accuracy depending on the pre-training task: sex classification gave better results than age prediction in accuracy for downstream tasks. One explanation for this can be the similarity in the design of the model and the loss function for sex classification and amyloid positivity classification, where both models perform binary classification (female/male and amyloid negative/positive respectively), with a binary cross-entropy loss function in the final layer. On the other hand, a regression model is used when pre-training for age prediction, where the activation function is linear, and the loss being minimized is MSE. This model is then converted to the classification model for the downstream task with the loss function changed to become binary cross-entropy. Another explanation may be the difference in subjects’ ages between ADNI and the UK Biobank with chronological age where the UK Biobank subjects are younger on average and may have less brain atrophy overall on the structural T1w MRI. As a result, the features learned during pre-training on age prediction, might not be as helpful in the downstream task.

**Figure 2.**
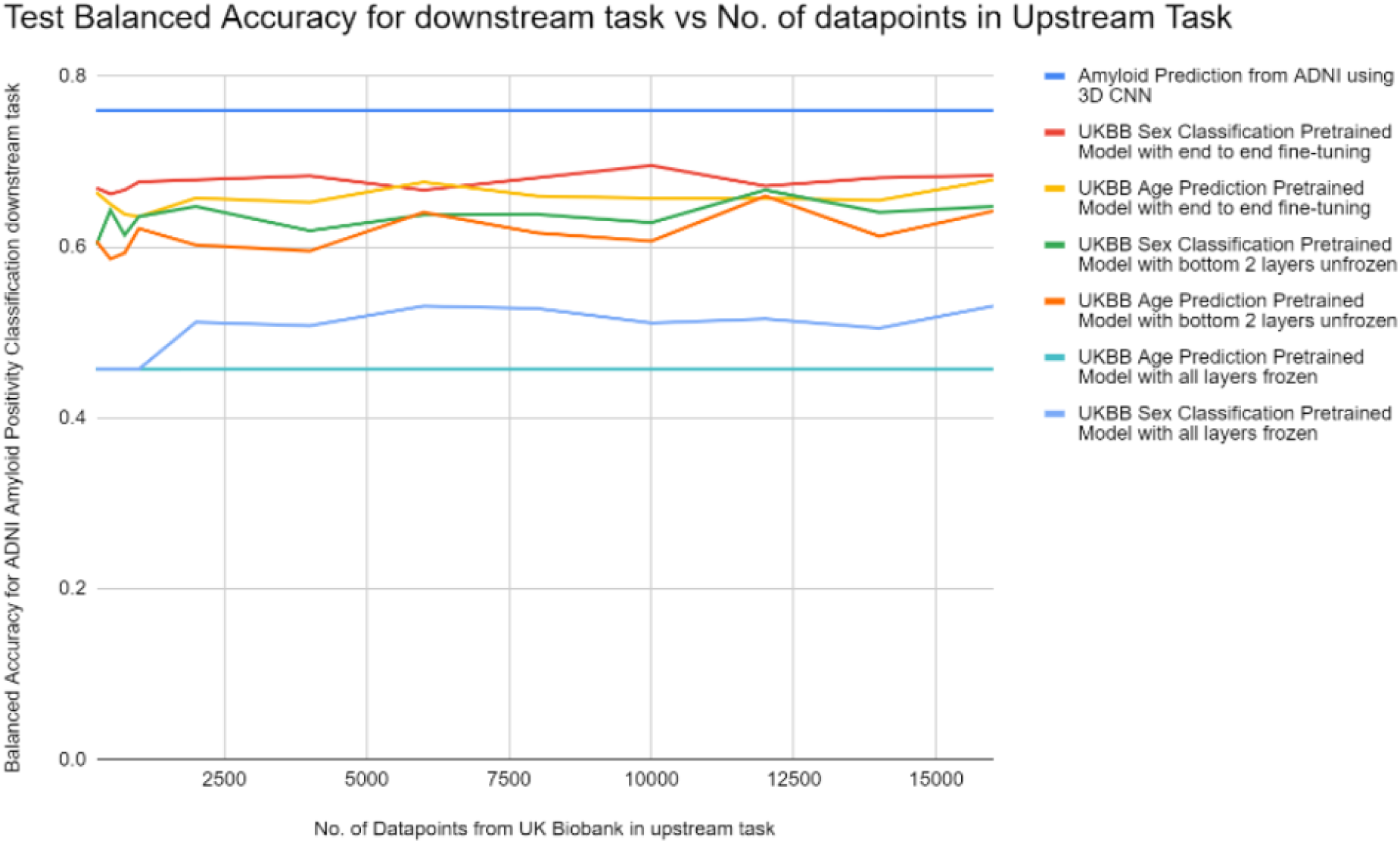
Plot of ADNI Test Set Balanced Accuracy vs % of training scans in pre-training from UK Biobank Data. The topmost line represents the balanced accuracy when the 3D CNN Model is trained from scratch on ADNI Data without any pre-training.

**Figure 3.**
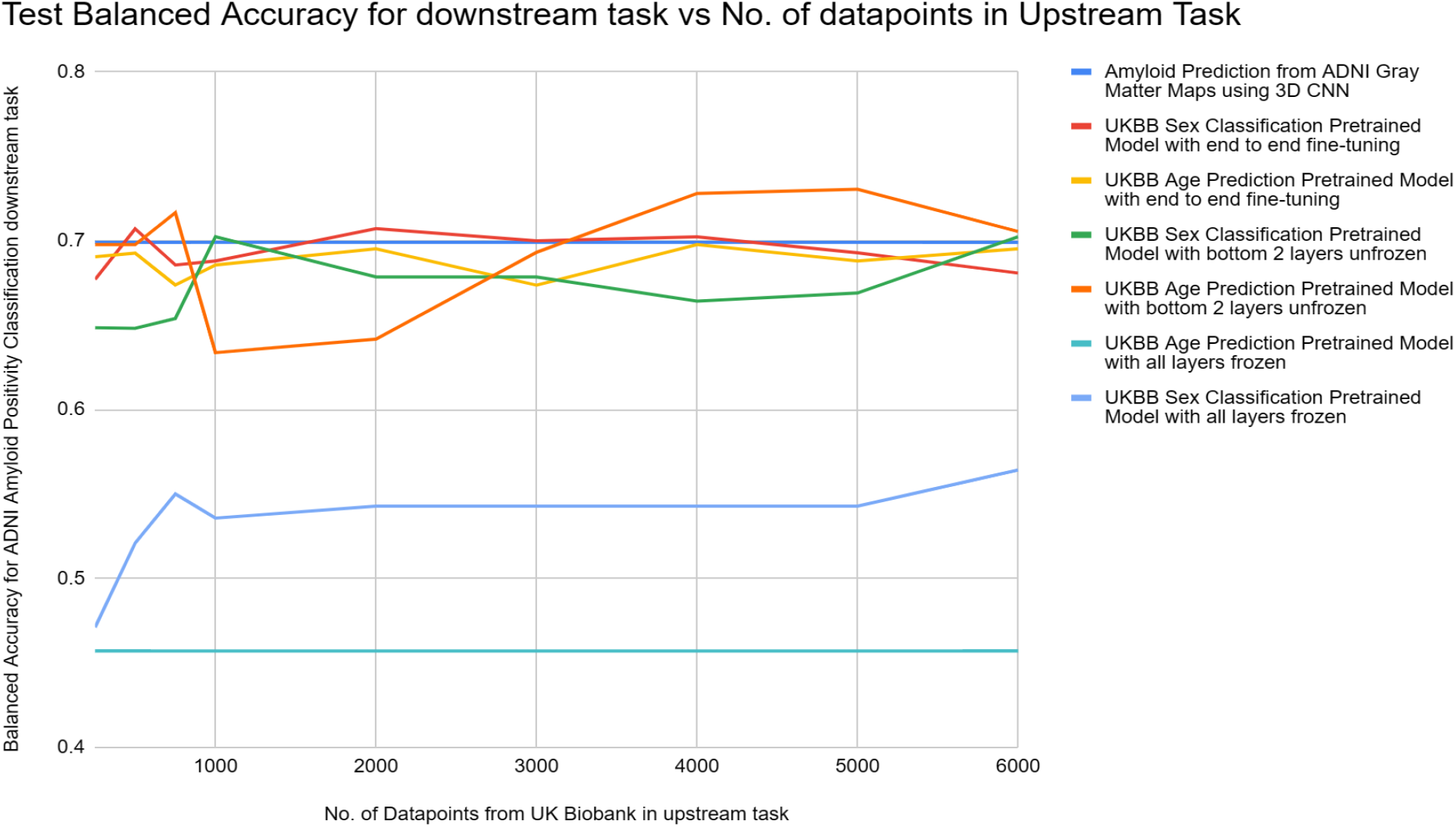
Plot of Gray Matter ADNI Test Set Balanced Accuracy vs % of training scans in pre-training from UK Biobank Data. Statistical testing revealed no evidence for performance gain with more pre-training data, at least in the range tested.

**Figure 4.**
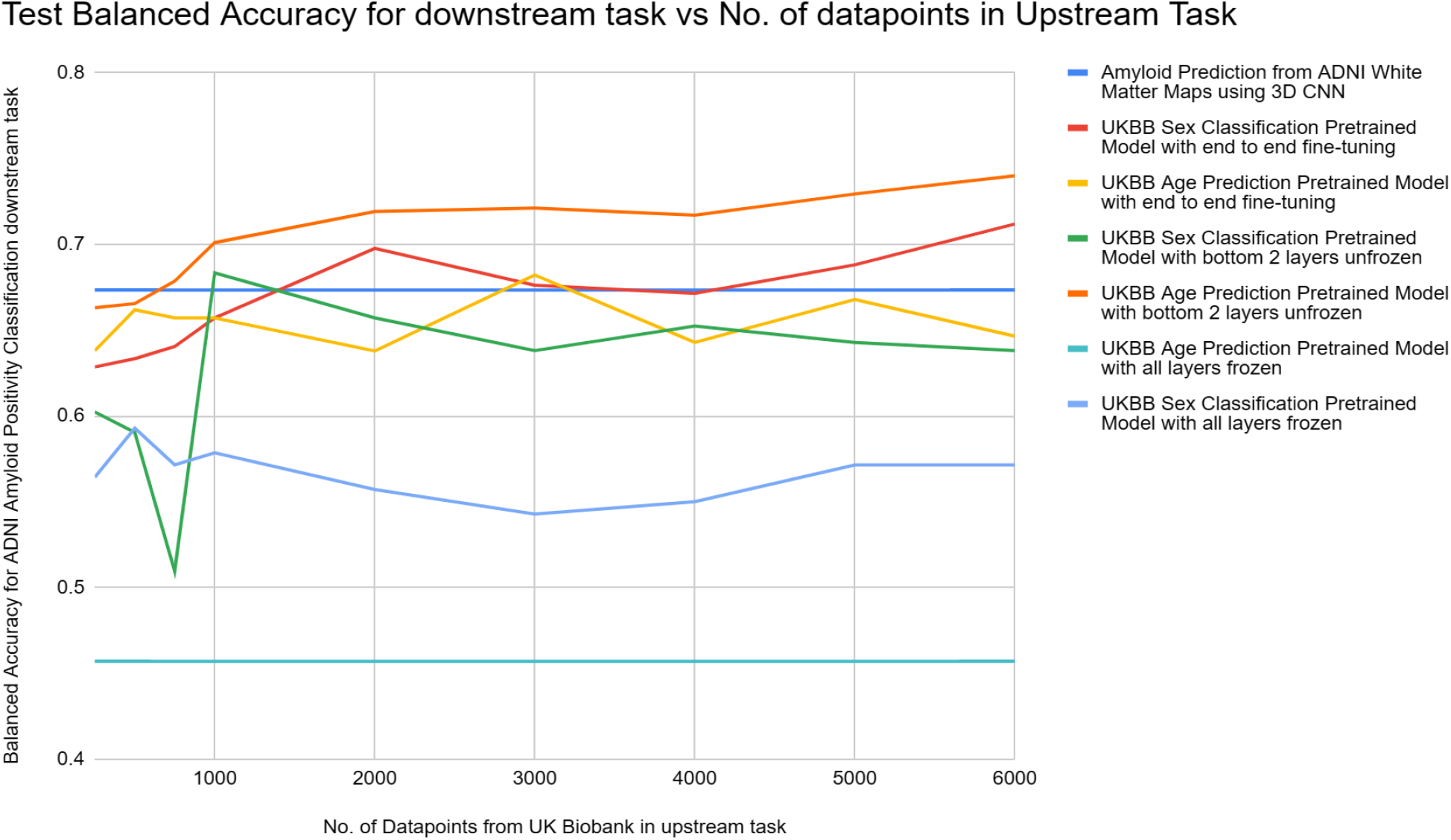
Plot of White Matter ADNI Test Set Balanced Accuracy vs % of training scans in pre-training from UK Biobank Data. Statistical testing revealed no evidence for performance gain with more pre-training data, at least in the range tested.

**Figure 5.**
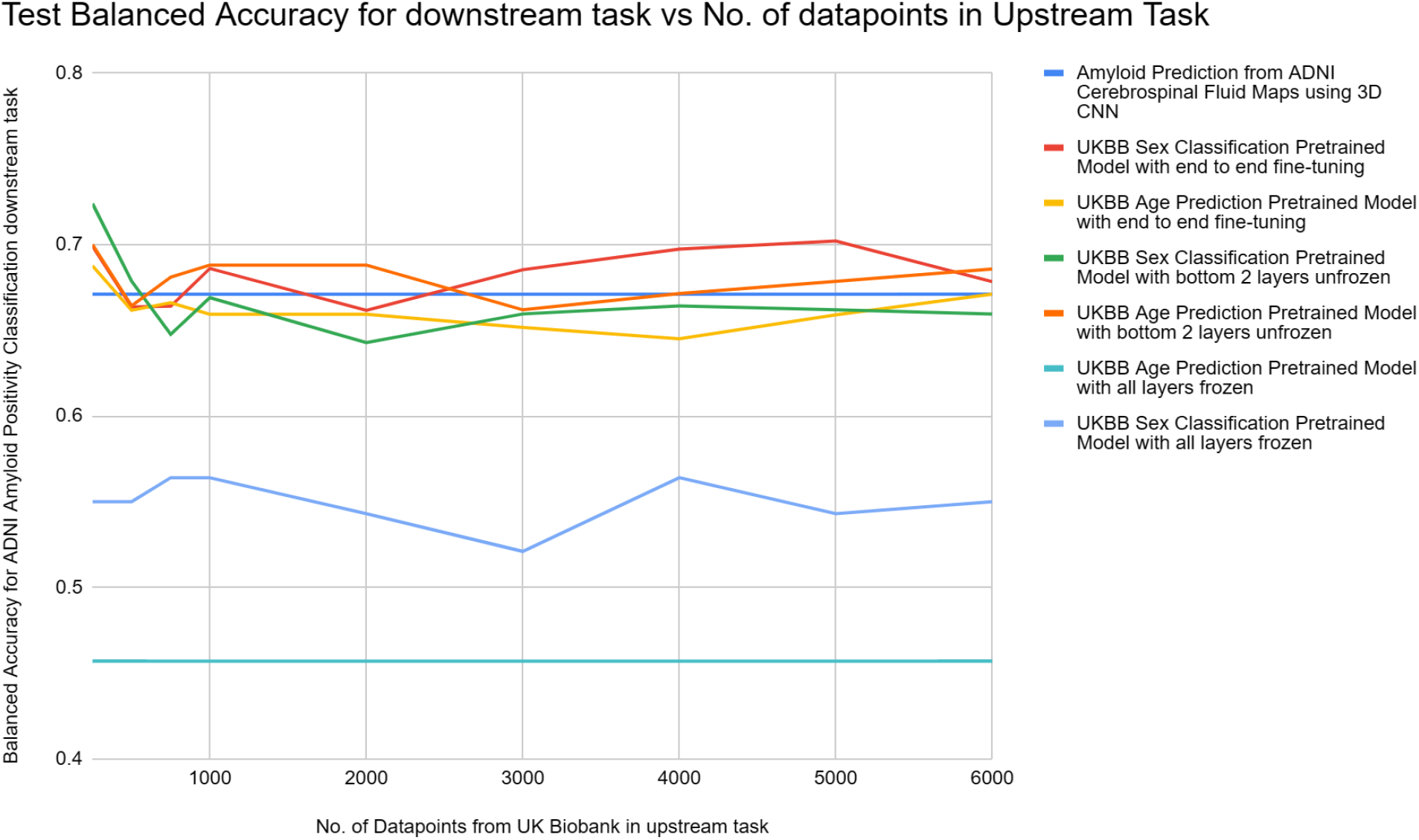
Plot of Cerebrospinal Fluid ADNI Test Set Balanced Accuracy vs % of training scans in pre-training from UK Biobank Data. Statistical testing revealed no evidence for performance gain with more pre-training data, at least in the range tested.

Pre-training helped in boosting the balanced accuracy in the downstream task in the case of white matter and to a small extent for CSF. The choice of pre-training task may also be important: as a pre-training task, age prediction gave better results than sex classification in terms of accuracy for the downstream tasks, except for the case when using CSF maps. Amyloid classification is typically based on other data sources such as amyloid- or tau-sensitive PET, or CSF biomarkers, which are all more invasive than structural brain MRI. A T1w MRI based model using derived GM, WM and CSF data is unlikely to be used in isolation, without the other data sources, but it can be fruitful for benchmarking, as T1ws are typically more widely available and cheaper to obtain compared to an amyloid PET scan. Thus, classifying amyloid positivity from T1ws and segmented GM, WM and CSF from T1w MRIs can form an important baseline based on which subjects might be selected for further, more intensive testing.

## VI. FUTURE WORK

Future work will use more paired training data, along with other data modalities such as FLAIR or diffusion MRI, for this challenging task. We will examine other deep learning architectures such as contrastive learning, or other CNN variants such as DenseNet-121, which has more depth than a standard 3D CNN. We would also like to extend pre-training techniques to additional patient datasets such as OASIS or from the AD Sequencing Project (ADSP).

## Acknowledgments

We thank the ADNI investigators and their public and private funders for creating and publicly disseminating the ADNI dataset. This research was supported by NIH grants R01AG058854, U01AG068057 and RF1AG057892.

